# A Reproducible Protocol for Embryonic Chicken Dorsal Root Ganglion Explant Culture and Quantitative Neurite Outgrowth Analysis

**DOI:** 10.64898/2026.09.14.745495

**Authors:** Yingcui Li, Alison Salinas, Lauren Wehner, Anna Keith, Yu Lok Evan Lo, Sharon Jennings, Raymond Xue, Robert S. Pijewski, Solaleh Miar, Lakshmi S. Nair, Kevin W.-H. Lo

## Abstract

**Background:** Primary embryonic dorsal root ganglion (DRG) explants offer an accessible and physiologically relevant organotypic *in vitro assay* platform that preserves native three-dimensional cellular interactions and supports robust neurite outgrowth, enabling studies of sensory neuron development and neurite outgrowth, as well as controlled assessment of neurotoxicity, and pharmacological compound effects.

**Methods:** Here we provide a comprehensive step-by-step method to micro-dissect and culture embryonic chicken DRGs and quantitatively measure neurite outgrowth. The protocol covers embryo preparation, vertebral column exposure, bilateral DRG microdissection, explant culturing, longitudinal phase-contrast imaging, and neurite image analysis.

**Results:** DRG explants maintained their overall ganglion morphology and remained attached to the culture surface throughout the five-day culture period. Untreated control explants exhibited limited neurite outgrowth, whereas nerve growth factor (NGF) treated explants developed progressively longer and denser radial neurite networks. Neurite outgrowth was readily visualized by live phase contrast microscopy and quantified using an Image-based neurite tracing method, demonstrating the responsiveness of the explant model to NGF and its suitability for quantitative assessment of neurite outgrowth.

**Conclusion:** This simple, low-cost protocol provides an *in vitro* assay platform for neurotoxicity, neuroregeneration, drug screening, and biomaterial evaluation.

## Introduction

Dorsal root ganglions (DRGs) are peripheral nervous system structures comprised of populations of sensory neurons cell bodies and satellite glial cells that relay signals from the periphery to the Central Nervous System (CNS) ^[1][2]^. During embryonic development, DRG neuron survival, differentiation, and neurite extension are highly regulated and dependent on neurotrophic factors like nerve growth factor (NGF) ^[3]^. Since perturbations to these events can lead to improper development of the sensory nervous system, DRGs during embryogenesis have been utilized as experimental platforms to examine neuronal growth and development, developmental neurotoxicology, and general responsiveness to bioactive molecules ^[4][5]^. The chicken embryo is advantageous for these studies due to its low cost, rapid development, ease of access for dissection, and availability of both left and right paired DRGs from individual embryos ^[6][7]^.

Primary DRG cultures can be grown as either dissociated cells or intact explants ^[8]^. Dissociated cultures allow analysis of cells at the single-cell level ^[9]^; however, this process involves enzymatic digestion and mechanical dissociation of tissue which can limit viability and alter neuron–glia relationships ^[10]^. While cell cultures derived from explants lack the benefit of single cell-level analysis, they maintain the intrinsic organization of the ganglion and enable longitudinal assessment of neurite outgrowth directly from a specific core of tissue ^[11]^. Although embryonic chicken DRG explants have been widely used, published methodologies differ in embryo stage, culture conditions, imaging schedules, and quantitative analysis, with many experimental details only briefly described ^[12]^. This variability limits reproducibility and broader adoption of the assay across laboratories. Previous studies performed on embryonic chicken DRGs investigated trypsin-mediated dissociation s^[13]^; however they demonstrated that cell yield and viability differed based on enzyme concentration and duration of exposure suggesting that additional optimization is necessary before dissociation can be implemented as a reliably reproducible assay ^[14]^. Due to these drawbacks, intact DRG explants provide an effective platform for assessing neurite outgrowth and treatment-dependent alterations in neuronal development ^[11]^.

Many previous studies have utilized chicken embryos at varying stages or cultured under different conditions ^[15][16]^. Additionally, methods are often briefly summarized in studies primarily focused on applications. Therefore, having a single standardized protocol outlining embryo preparation, DRG isolation, explant culturing, longitudinal imaging, and quantitative analysis would increase reproducibility and adoption of this model.

Here, we provide here a detailed protocol for isolation, culture and quantification of embryonic chicken DRG explants. The method includes incubation of fertilized eggs, embryo preparation, exposing of the vertebral column, bilateral microdissection of DRGs, explant culture and longitudinal phase-contrast microscopy for quantification of neurite outgrowth. Mean neurite length is used as a quantitative phenotypic endpoint, and NGF-induced neurite outgrowth serves as a positive-control response to demonstrate assay performance. Overall, this assay represents a simple, inexpensive and reproducible primary neuronal assay that can be implemented for developmental neurotoxicity testing, phenotypic drug screening, peripheral nerve regeneration assays, and testing of biomaterials that enhance or inhibit sensory neurite outgrowth.

### Protocol 1. Embryo Preparation

#### Materials

- Fertilized White Leghorn chicken eggs
- Egg incubator (37.5°C, 50–60% humidity)
- 70% ethanol
- Sterile 60-mm Petri dishes
- Hank’s Balanced Salt Solution (HBSS)
- Fine forceps (disinfected with 70% ethanol)
- Spring scissors (disinfected with 70% ethanol)
- Transfer pipette
- Stereomicroscope (Dissecting)

#### Procedure

1. Incubate fertilized chicken eggs at 37.5°C with 50–60% relative humidity. The day eggs are placed in the incubator is designated embryonic day 0 (E0); embryos are collected on E11.
2. Remove the desired number of eggs from the incubator and allow them to cool to room temperature for approximately 2–3 min.
3. Wipe the eggshell thoroughly with 70% ethanol and allow the surface to air dry.
4. Crack the eggshell carefully around its equator using sterile forceps or scissors.
5. Open the shell and gently transfer the intact embryo into a sterile 60-mm Petri dish containing 2–3 mL cold sterile HBSS.
6. Remove excess egg albumen and yolk by gently rinsing the embryo with fresh HBSS.
7. Decapitate the embryo using fine spring scissors to euthanize it.
8. Transfer the embryo into a clean 60-mm Petri dish containing approximately 2-3 mL fresh cold HBSS.
9. Position the embryo ventral side upward under a stereomicroscope.
10. Remove the ventral organs using fine forceps to expose the vertebral column.
11. Remove the forelimbs, hindlimbs, skin, and surrounding musculature while preserving the integrity of the spinal column.
12. Proceed immediately to DRG microdissection.

### Protocol 2. Microdissection of Embryonic Chicken Dorsal Root Ganglia (DRGs)

#### Materials

- Sterile HBSS
- Dumont #5 forceps (2 pairs) (disinfected with 70% ethanol)
- Fine spring scissors (disinfected with 70% ethanol)
- 60-mm sterile Petri dishes
- Fire-polished Pasteur pipette (or wide-bore transfer pipette)
- Stereomicroscope
- 70% ethanol

#### Procedure

1. Disinfect all dissection instruments by soaking them in 70% ethanol for at least 15 minutes and allow them to air dry under sterile conditions.
2. Transfer the excised vertebral column prepared in Protocol 1 into a fresh Petri dish containing cold sterile HBSS.
3. Orient the vertebral column dorsal side upward under the stereomicroscope.
4. Stabilize the vertebral column with one pair of Dumont #5 forceps.
5. Using a second pair of forceps, gently separate the vertebral arches to expose the spinal cord.
6. Carefully lift and remove the spinal cord, exposing the paired DRGs along both sides of the vertebral column.
7. Identify each DRG as a small translucent oval structure attached to the dorsal nerve roots.
8. Beginning at the cervical region and progressing caudally, gently detaching each DRG by grasping the adjacent connective tissue or nerve root. Avoid directly compressing the ganglion body.
9. Remove excess connective tissue while preserving the overall architecture of the ganglion.
10. Pre-wet the fire-polished Pasteur pipette or wide-bore transfer pipette with sterile HBSS before collecting DRGs to minimize tissue adherence to the pipette wall.
11. Transfer each isolated DRG immediately into culture medium using a fire-polished Pasteur pipette or wide-bore transfer pipette.
12. Repeat the procedure until all desired DRGs have been collected (typically 10– 12 DRGs per culture dish).
13. Proceed immediately to explant plating to maintain tissue viability.

### Protocol 3: DRG Explant Culture

#### Materials

- Dulbecco’s Modified Eagle Medium/Nutrient Mixture F-12 (DMEM/F-12)
- Fetal bovine serum (FBS)
- Penicillin–streptomycin (100 U/mL penicillin, 100 μg/mL streptomycin)
- Sterile 35-mm tissue culture dishes
- Fire-polished Pasteur pipette or wide-bore pipette tip
- Humidified CO₂ incubator (37°C, 5% CO₂)
- Phase-contrast microscope
- Recombinant nerve growth factor (NGF)

#### Method

1. Prepare complete culture medium consisting of DMEM/F-12 supplemented with 10% fetal bovine serum and 1% penicillin–streptomycin.
2. Add 2 mL of complete culture medium to each sterile 35-mm tissue culture dish.
3. Transfer freshly isolated DRG explants individually into the culture dish using a fire-polished Pasteur pipette or wide-bore pipette tip.
4. Arrange the explants evenly across the culture dish with sufficient spacing to prevent neurite overlap during subsequent growth. Typically, 10–12 DRGs are plated per 35-mm dish.
5. Allow the explants to settle naturally onto the culture surface before moving the culture dish.
6. Incubate the cultures at 37°C in a humidified atmosphere containing 5% CO₂.
7. NGF treatment: Following explant attachment, replace the culture medium with complete medium containing 60 ng/mL nerve growth factor (NGF). Maintain untreated explants in complete medium without NGF as the negative control. Replace the treatment medium according to the experimental schedule.
8. Maintain the cultures under standard conditions throughout the experimental period. For extended culture, carefully remove 1 mL of medium and replace it with 1 mL of fresh prewarmed complete medium.
9. Acquire phase-contrast images at predetermined time points using identical microscope settings for all samples.

### Protocol 4: Phase-Contrast Microscopy and Neurite Outgrowth Quantification

#### Materials

- Inverted phase-contrast microscope equipped with a digital camera
- Image acquisition software
- Computer with ImageJ/Fiji software
- Simple Neurite Tracer plugin
- Calibrated stage micrometer

#### Method

1. Remove the DRG culture dish from the incubator at the designated imaging time point.
2. Place the culture dish on the stage of an inverted phase-contrast microscope.
3. Locate an individual DRG explant under low magnification.
4. Center the DRG explant and its surrounding neurite field within the microscope view.
5. Acquire a phase-contrast image of the DRG explant.
6. Repeat the imaging procedure for each DRG explant in the culture dish.
7. Use the same microscope magnification and imaging conditions for all DRGs and experimental groups.
8. Save the acquired images for neurite outgrowth analysis.
9. Open each DRG image in ImageJ.
10. Select Plugins → Segmentation → Simple Neurite Tracer.
11. Use the neurite-tracing tool to identify and trace neurites extending outward from the DRG explant.
12. For each DRG, identify the five longest neurites.
13. Trace each selected neurite from the edge of the DRG body to its distal endpoint.
14. Record the length of each of the five neurites.
15. Calculate the average length of the five measured neurites.
16. Record this average as the mean neurite length for that DRG explant.
17. Repeat the measurement procedure for all DRG explants and experimental groups.
18. Use the mean neurite length for each DRG for subsequent comparison and statistical analysis.

## Results

### Embryonic chicken DRG isolation and explant preparation

Figure 1 illustrates the sequential preparation and identification of embryonic chicken DRGs. Figure 1A summarizes the extraction procedure, while Figure 1B shows the prepared embryo with the vertebral region exposed. Bilateral DRGs were subsequently identified along the vertebral column (Figure 1C), and individual intact DRGs were microdissected for explant culture (Figure 1D).

**Figure 1.**
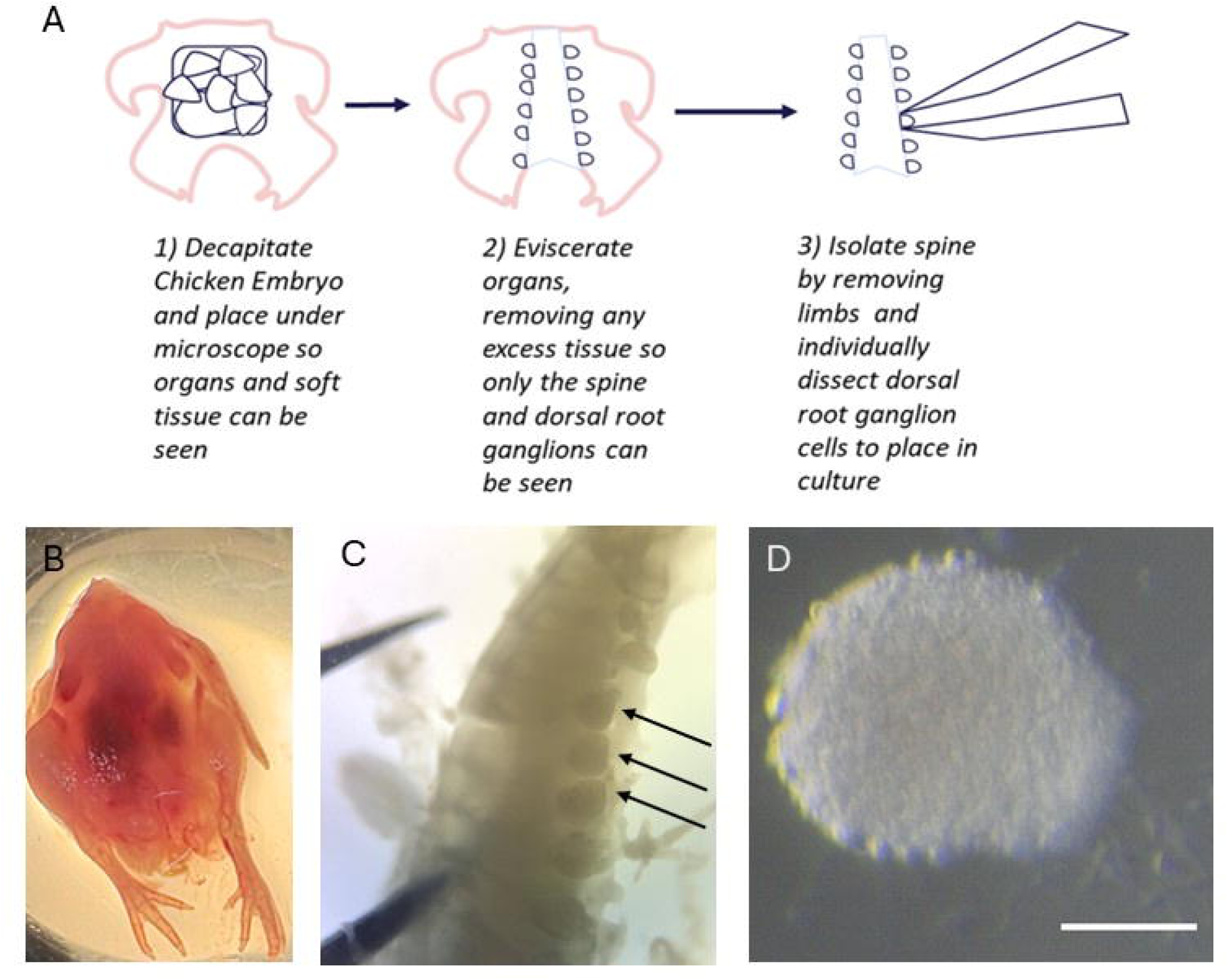
Embryonic chicken anatomy and dorsal root ganglion (DRG) isolation. (A) Schematic diagram for embryonic chicken dorsal root ganglion extraction. The embryo is first decapitated and positioned ventral side up for dissection. The ventral organs and surrounding soft tissues are then removed to expose the vertebral column and bilateral dorsal root ganglia. Finally, the vertebral column is isolated, and individual DRGs are microdissected and transferred for explant culture. (B) Ventral view of the E11 chicken embryo prior to evisceration, with the abdominal organs still intact. (C) Higher-magnification view of the vertebral column showing bilateral DRGs (arrow). (D) Representative isolated DRG explant immediately after isolation and prior to culture. Scale bar = 200 μm.

Figure 2 summarizes the entire experimental workflow. Steps included egg incubation and embryo staging, embryo preparation, vertebral column exposure, bilateral DRG microdissection, explant plating and treatment, longitudinal phase-contrast microscopy, and Image-based neurite analysis. The experimental workflow allowed us to track DRG explants over the course of culture, beginning with their isolation and ending with quantitative measurement of neurite outgrowth.

**Figure 2.**
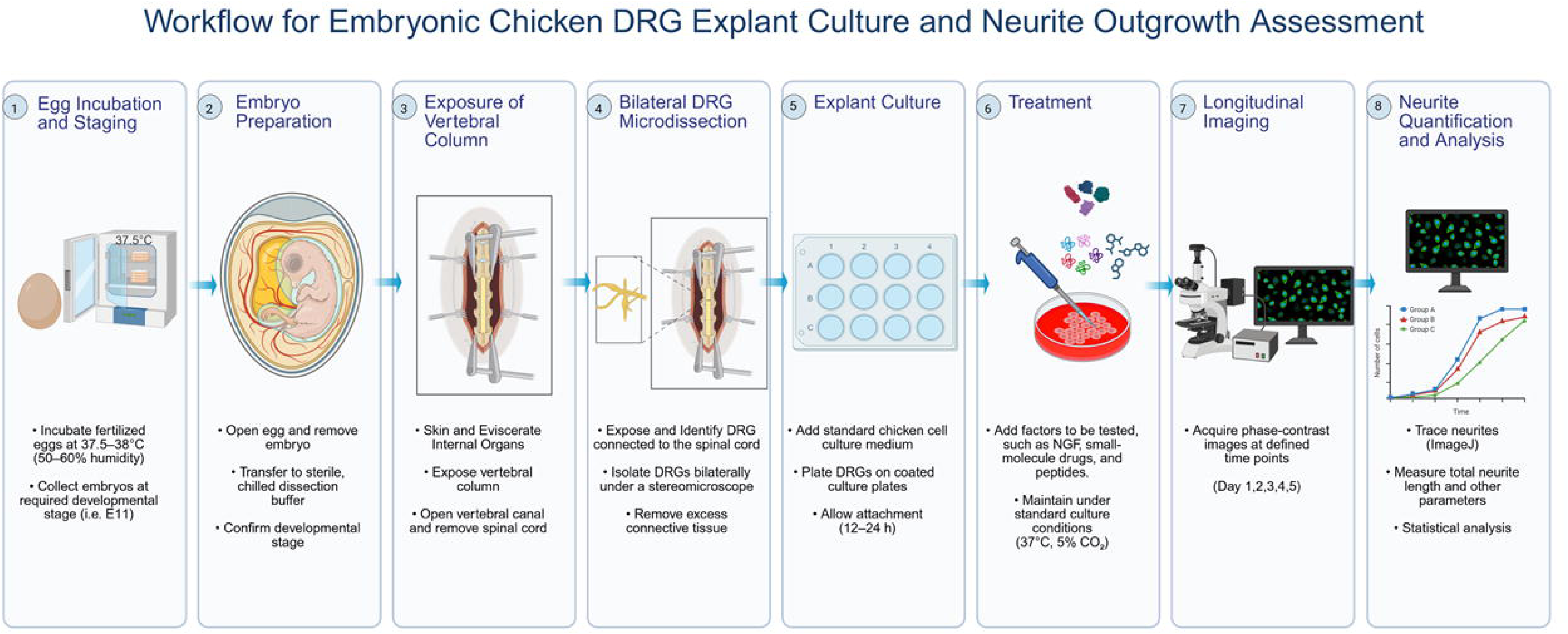
Overall workflow for the isolation of embryonic chicken dorsal root ganglia and explant preparation. The graphic depicts stepwise processes used for preparation of embryonic chicken tissue explants used for DRG isolation. The embryo is decapitated and eviscerated, and then the limbs, skin, musculature and surrounding connective tissue are removed to expose and isolate the vertebral column. The spinal cord is removed exposing the bilateral DRGs that are individually dissected and collected for explant culture (Image was generated by BioRender).

### NGF promoted neurite outgrowth over five days

NGF served as a positive control as it promotes sensory neurite outgrowth ^[17]^, thus allowing us to ensure that our assay would be responsive. DRG explants were monitored for five days to allow visualization of neurite extension over time. Basal untreated control explants exhibited minimal neurite outgrowth and generally maintained a rounded morphology. By comparison, explants treated with NGF developed increasingly large neurite arbors with extensive radial outgrowth by Day 5 (Figure 3). This result illustrates that the above-described cultured embryonic chicken DRG remained attached and responsive to neurotrophic stimulation throughout the five-day assay period.

**Figure 3.**
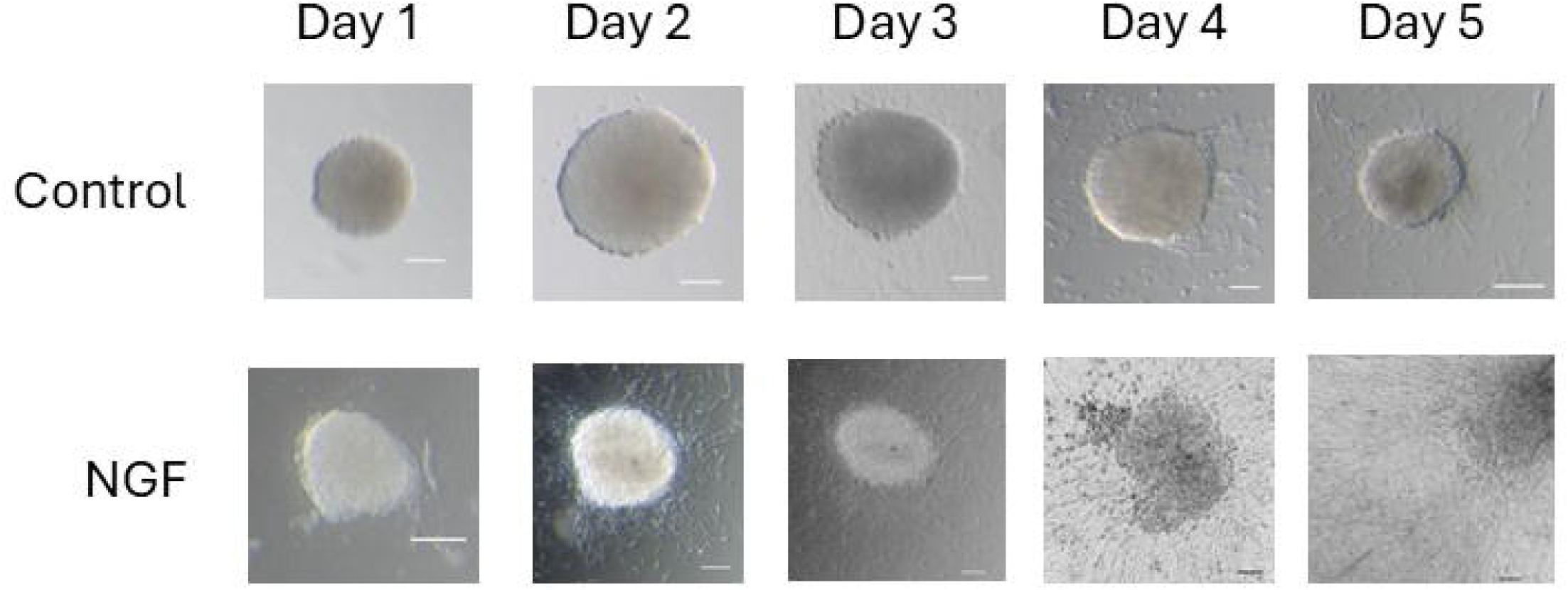
Effect of nerve growth factor on neurite outgrowth from embryonic chicken dorsal root ganglion explants. Representative phase-contrast images of DRG explants cultured for five days under control conditions without treatment (upper row) or with nerve growth factor treatment (lower row). Images show progressive explant attachment and neurite extension, with more pronounced neurite outgrowth observed in the NGF-treated group. Scale bar = 500 μm.

### Image-based quantification of neurite length

The neurite quantification methodology is depicted in Figure 4. Phase-contrast images were processed in ImageJ with the Simple Neurite Tracer plugin. Figure 4A shows a representative NGF-treated explant with extensive neurite extension, whereas the untreated control exhibits more limited outgrowth (Figure 4B). Individual neurites were traced starting at the perimeter of the DRG body and ending at the distal endpoint (Figure 4C). The five longest neurites per explant were measured and averaged to determine a mean neurite length for that explant. As shown by representative analysis, the NGF-treated group had longer mean neurite lengths than the untreated control group, corroborating the qualitative differences seen in longitudinal images (Figure 4D). Taken together, these results show that the protocol allows for reproducible DRG isolation, maintenance of explant cultures, visualization of treatment-dependent neurite outgrowth, and quantitative, image-based analyses.

**Figure 4.**
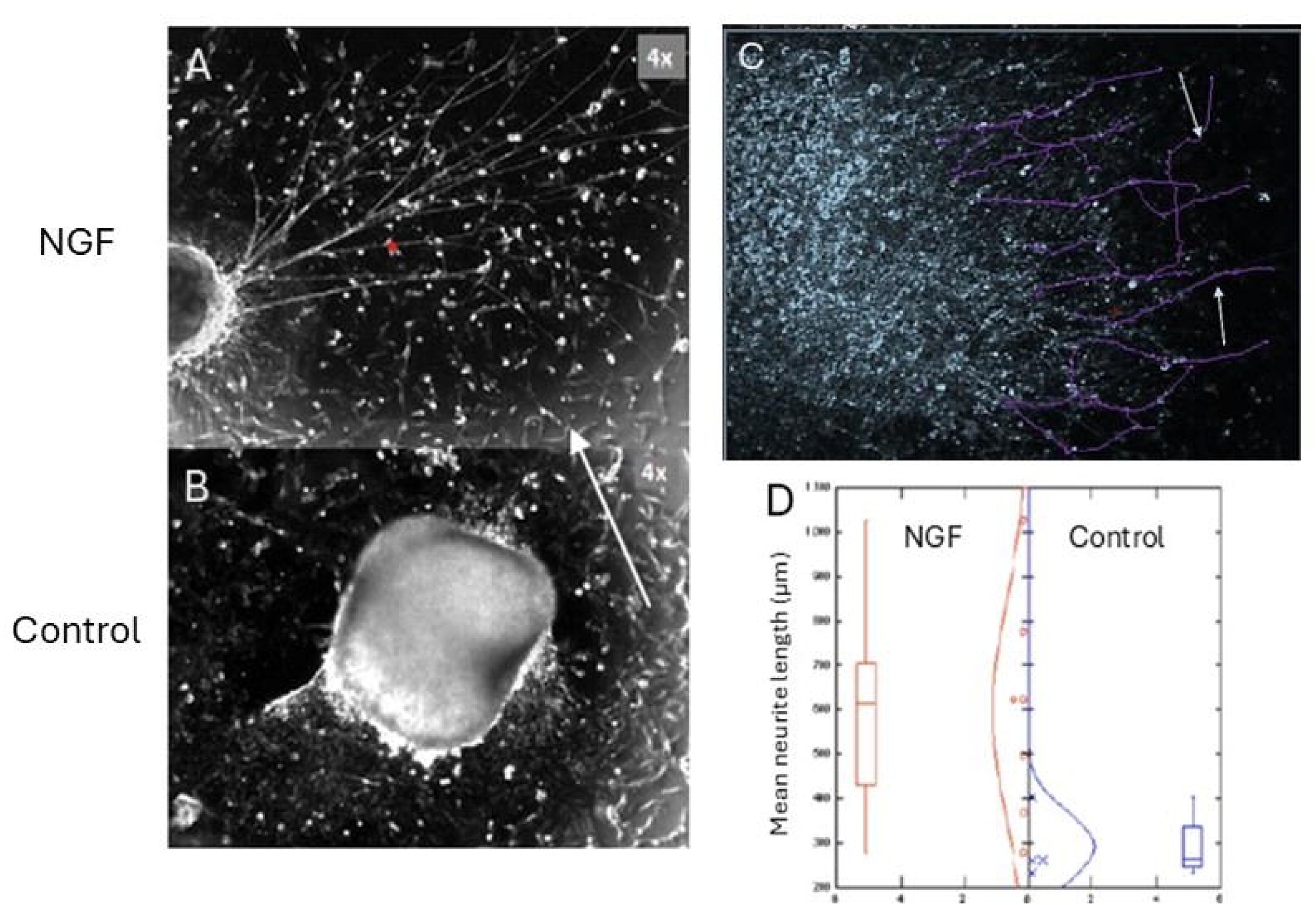
Image-based measurement of neurite outgrowth from embryonic chicken dorsal root ganglion explants. Representative phase-contrast images of DRG explants cultured in the presence of nerve growth factor (NGF) **(A)** or under untreated control conditions **(B)**. **(C)** Representative ImageJ analysis showing neurites traced from the edge of the DRG explant to their distal endpoints using the Simple Neurite Tracer plugin. **(D)** Quantification of mean neurite length in NGF-treated and untreated control DRG explants. For each explant, the five longest measurable neurites were traced and averaged to obtain a mean neurite length. Also note that neurite tracing was performed on calibrated phase-contrast images using ImageJ; scale information is provided in the corresponding representative microscopy images.

## Discussion

Here we describe a detailed protocol for isolation, culture, and quantification of neurites from embryonic chicken dorsal root ganglion (DRG) explants. By combining micro dissection, explant culture, longitudinal imaging, and image-based neurite quantification, this method offers a reproducible platform for assessing sensory neuron development and treatment-induced changes in neurite growth.

Key steps for successful execution of this protocol include gentle tissue handling, stable explant attachment, consistent imaging, and standardized neurite tracing. A summary of common technical pitfalls and possible troubleshooting strategies are listed in Table 1. These suggestions should aid users in identifying factors that introduce variability and standardizing DRG isolation, culturing, and neurite outgrowth measurements. Although mean neurite length serves as a primary readout in this protocol, additional assays such as neurite number, branching, outgrowth area, explant attachment, cell viability, or molecular markers may also be incorporated depending on the purpose of the assay^[18][11][19]^.

**Table 1.** Potential challenges and troubleshooting.

| Potential challenge | Possible cause | Troubleshooting |
| --- | --- | --- |
| Embryo or culture contamination | Inadequate sterilization of eggs, instruments, or work area | Wipe eggs with 70% ethanol, disinfect instruments before dissection, and maintain sterile technique throughout isolation and culture. |
| Vertebral column damaged during preparation | Excessive pulling or cutting during removal of organs, limbs, and surrounding tissues | Remove tissues gradually while preserving the integrity of the vertebral column. |
| DRGs identification | Incomplete removal of the spinal cord or surrounding connective tissue | Carefully open the vertebral arches and remove the spinal cord to expose the bilateral DRGs. |
| DRG damaging during isolation | Direct compression of the ganglion with forceps | Grasp the adjacent nerve root or connective tissue rather than the DRG body. |
| Explant attachment in culture dish | Culture dish moved immediately after plating or insufficient settling time | Allow DRGs to settle naturally before moving the dish and minimize disturbance during early culture. |
| DRGs detach during medium changes | Medium added or removed too rapidly | Perform medium changes slowly and image cultures before medium replacement. The thesis specifically notes that careful handling is needed to prevent loss or detachment of DRGs. |
| Neurites overlap between explants | DRGs plated too closely together | Space DRGs evenly across the culture dish and reduce the number of explants if necessary. |
| Limited neurite outgrowth | Damaged explant, poor attachment, or absence of an appropriate growth stimulus | Use intact, well-attached DRGs and include an NGF-treated positive-control group to confirm assay performance. |
| Poor or inconsistent images | Differences in magnification, illumination, or focus | Use the same microscope settings and magnification for all samples and time points. |
| Fewer than five measurable neurites | Weak neurite growth or damaged explant | Exclude DRGs that do not produce at least five clearly measurable neurites, consistent with the thesis analysis method. |
| Variable neurite measurements | Inconsistent selection or tracing of neurites | Measure the five longest neurites using the same ImageJ tracing criteria and average the measurements for each DRG. |

Compared to mouse embryonic DRG cultures, embryonic chicken DRGs allow for a simplified, low-cost, and more readily accessible ex vivo model system for neurite outgrowth studies^[20]^. Fertilized eggs are abundant and each embryo produces multiple paired DRGs allowing for greater throughput of experiments with less expense and variability. While mouse DRGs will always be useful for transgenic and disease specific investigations^[21]^, embryonic chicken DRGs provide an effective model system to study developmental neurobiology, neurotoxicity, drug screening, and biomaterial assessment.

### Conclusion and Perspective

An advantage of the embryonic chicken DRG explant assay is that it is an easy-to-use *ex vivo* system that can be readily applied to phenotypic drug discovery assays, developmental neurotoxicity assays, and assays to study regenerative medicine mechanisms. Additional automation through incorporation with high-content imaging and AI-based neurite tracing, microfluidic culture devices, three-dimensional culture matrices, and molecular or electrophysiological endpoints would increase both assay throughput and assay physiological relevance. Further, since this assay allows for rapid quantification of neurite outgrowth responses, it may serve as an early screen for regenerative medicine therapeutics before advancing into more sophisticated *in vivo* studies.

## Ethics Statement

All procedures involving chicken embryos were conducted in accordance with institutional guidelines. Fertilized chicken embryos were used only through embryonic day 11 (E11), a developmental stage that does not require Institutional Animal Care and Use Committee (IACUC) approval under U.S. federal regulations.

## Disclosure

This manuscript was polished for language and clarity using AI-based tool (ChatGPT5.4). The authors reviewed and took full responsibility for the content.

